# Re-evaluating evolution in the HIV reservoir

**DOI:** 10.1101/140731

**Authors:** Daniel I. S. Rosenbloom, Alison L. Hill, Sarah B. Laskey, Robert F. Siliciano

**Author notes:** Address correspondence to Robert Siliciano.

## Abstract

Despite antiretroviral therapy (ART), a latent reservoir of replication-competent HIV-1 persists in resting memory CD4^+^ T-cells and precludes cure^1-6^. Lorenzo-Redondo *et al*.^7^ analyzed HIV-1 sequences collected from three individuals during the first six months of ART, discovered specific patterns of sequence evolution, and concluded that viral replication persists during therapy. We believe these evolutionary patterns are artifacts of rapidly decaying viral subpopulations present during the first months of therapy and are not characteristic of the long-lived reservoir. The study therefore provides no evidence that ongoing replication is an additional barrier to cure for treated individuals who consistently maintain low viral loads.

---

Lorenzo-Redondo *et al*. collected samples before and three and six months after treatment initiation, when labile viral populations dominate and change rapidly. Prior to treatment, most HIV-1 DNA in resting CD4^+^ T-cells exists in an unintegrated state decaying with a half-life of days^8,9^. Another major population of infected resting cells decays with a half-life of weeks^10^. The latent reservoir of integrated proviruses, observed in blood and lymphoid tissue^1^, is smaller and decays with half-life of ~4 years^5,6^. Lifelong persistence of this reservoir is determined by longevity and proliferation of the infected cells^11^. Initiation of ART blocks new infection from replenishing these populations, revealing their different lifespans. Differential decay causes dramatic shifts in infected cell populations in the first six months of ART, making suspect any conclusions about viral evolution gleaned from this period. Below, we support this claim by simulating differential decay and replicating the analysis of Lorenzo-Redondo *et al*. on the simulated data. We find that false signals of viral evolution – and ongoing viral replication – often appear.

To estimate size and decay of labile compartments, we examined a cohort of seven early-treated individuals, which we consider comparable to the two early-treated participants in the Lorenzo-Redondo *et al*. study. Blankson *et al*.^10^ used the quantitative viral outgrowth assay (qVOA) on blood samples to detect resting CD4^+^ T-cells harboring replication-competent HIV-1. At ART initiation, infected cell frequencies greatly exceeded those of individuals on long-term ART. A multi-log, multi-phasic decay over the first year of therapy reduced frequencies to levels typically observed during long-term ART. Fitting to the most extensively sampled individual, we inferred a large, fast-decaying population, a smaller population with slower decay, and a very small persistent reservoir, approximated as constant (Fig. 1). At zero, three, and six months following ART initiation, labile populations comprise 99.99%, 96.2%, and 76% of infected resting cells, respectively, masking the persistent reservoir. The RNA-based assays performed by Lorenzo-Redondo *et al*. on lymphoid tissue paint a similar picture: the infection decays rapidly over the first three to six months, eventually dwindling to a more stable state >3 orders of magnitude smaller than the pretreatment population (Lorenzo-Redondo *et al*., Extended Data Figure 1). Regardless of sequencing depth, the limited number of infected cells in a blood draw or tissue biopsy likely prevents the persistent reservoir from being sequenced at early timepoints. The genetic diversity of this reservoir only emerges later. Latency studies are therefore generally restricted to participants who have received suppressive ART for >6 months, a precaution not taken by Lorenzo-Redondo *et al*.

**Figure 1.**
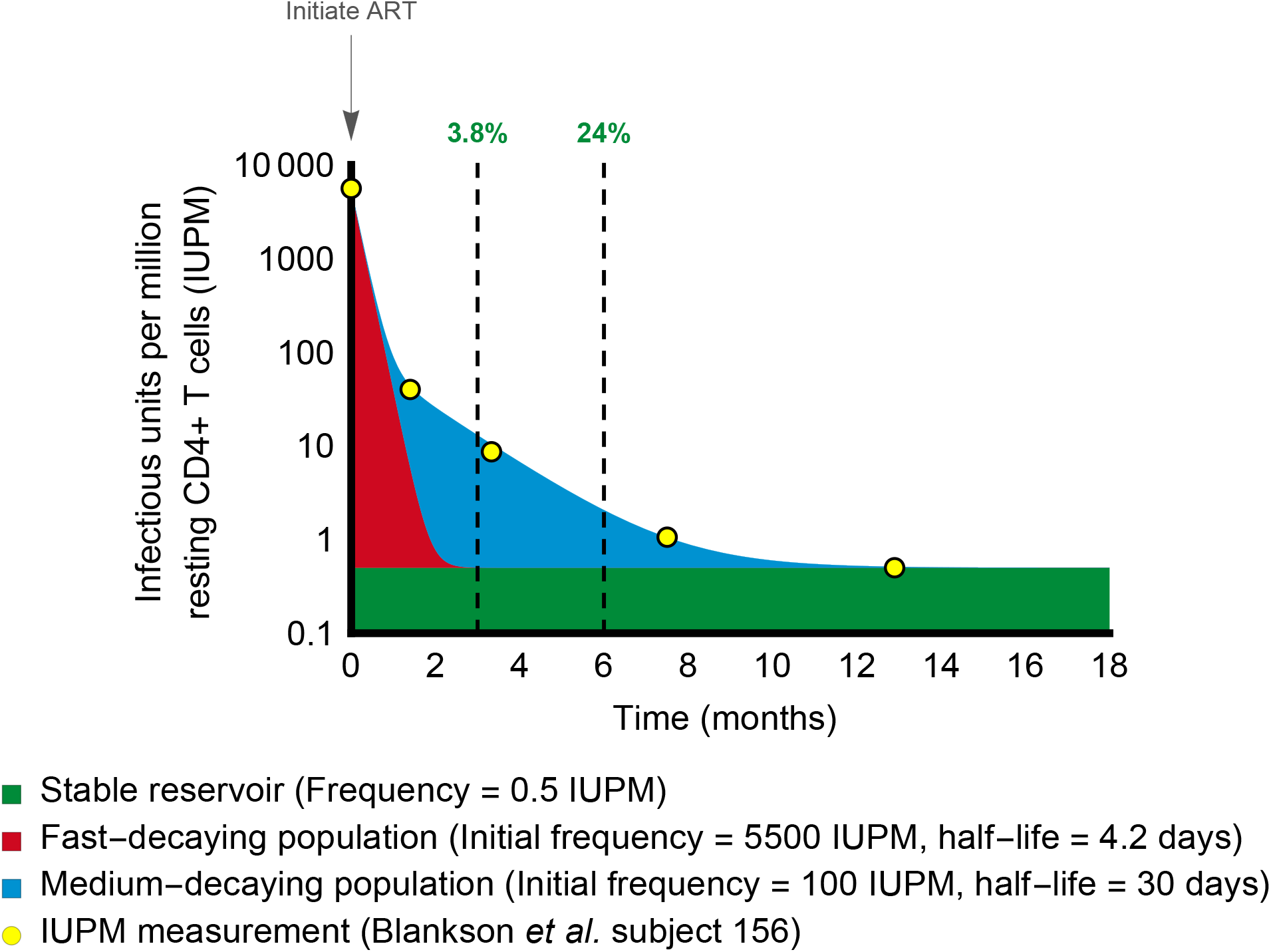
For the first six months of ART, the persistent latent reservoir makes up a minority of sampled sequences. Yellow dots show longitudinal measurements of inducible, replication-competent HIV-1 in resting CD4^+^ T cells from a patient studied by Blankson *et al*.^10^. Infection frequency was measured by qVOA and is reported as infectious units per million cells (IUPM). Intact, unintegrated HIV-1 DNA in recently infected cells can be detected in this assay because cellular activation stimulates completion of the viral life cycle^1,8,10^. The solid-colored regions show estimated sizes of the underlying populations that combine to yield the observed tri-phasic decay of IUPM. A latent reservoir (green, proportion shown at 3 and 6 months) is established before treatment and persists despite ART. This reservoir decays at a very slow rate^5,6^ that can be approximated as constant over the one-year period represented here. Initially, there is a large, rapidly decaying viral population (red) likely to include unintegrated viral genomes. A smaller population of infected cells decays at a moderate rate (blue). Consequently, at therapy initiation, <0.01% of resting CD4^+^ T cells with replication-competent virus belong to the stable reservoir, while 98% belong to the large, fast-decaying population. Only after a year of therapy would the stable reservoir exceed 95%. All patients in the study showed similar decay in which the proportion of infected cells belonging to the latent reservoir did not stabilize within the first six months of therapy^10^. Note that cells with infectious provirus are generally outnumbered by orders of magnitude by cells containing defective provirus. The percentages given here therefore overestimate the proportion of all sampled HIV-1 DNA that represents the persistent latent reservoir of replication-competent virus.

Brodin *et al*.^12^ suggested that decay of labile populations may produce false signals of evolution during treatment, even in the absence of viral replication. We used computer simulations of viral populations during acute infection and treatment to confirm this hypothesis. Simulated virus replicated and seeded subpopulations for four months, and treatment then blocked replication for six months. During treatment, labile subpopulations decayed, while a stable reservoir persisted, as in Fig. 1. Nearly 12,000 simulations were subjected to the tests performed by Lorenzo-Redondo *et al:* genetic divergence from start to end of therapy, evolutionary rate calculations, and measurement of clock-like signal in maximum-likelihood trees (Supplementary Tables 1 and 2).

We tested a range of parameters defining growth and competition in the pre-treatment viral population and selected 8,000 simulations with realistic viral diversity. Depending on parameter values, up to 57% of simulations produced a false impression of clock-like evolution according to all three tests used by Lorenzo-Redondo *et al*. (Supplementary Methods). For comparison, of the 17 gene/tissue combinations studied by the authors, 11 (65%) produced a signature of evolution according to the two tests for which statistics were explicitly presented (Lorenzo-Redondo *et al*., Extended Data Tables 1 & 2). Decay of labile compartments can remove positively selected variants that occurred before treatment, revealing ancestral genotypes – which masquerade as the product of new mutation during treatment. Strong positive selection (anywhere in the genome, not only in the sequenced region) is required to generate a false impression of evolution. We believe that this mechanism is realistic, as rapid selective sweeps, caused by CTL escape mutations with selective coefficients of 20% or more, typify acute infection^13^.

To support our argument with actual sequence data and without assuming a selection coefficient, we simulated reservoir seeding and post-treatment decay using HIV-1 *gag* sequences obtained from three individuals during untreated acute infection^14^. Appearance of post-treatment “evolution” again depended on pre-treatment dynamics. The individual with most extensive sequence changes, suggesting strong pre-treatment selection, passed a phylogenetic test of forward evolution in 92 of 100 replicate simulations (Supplementary Methods, Supplementary Table 3, Extended Data Table 1).

Fig. 2 illustrates how decay of labile compartments produces the appearance of evolution in an example simulation (Replicate 91 of Parameter Set 48, Supplementary Table 2). Before treatment, the population diverges through mutation and selection. When treatment starts, the most diverged sequences die out, resulting in a decline in divergence. If divergence is measured not from the origin of infection, but rather (as by Lorenzo-Redondo *et al.)* from the most common genotype at treatment initiation, this retreat towards the origin is misinterpreted as increasing divergence (Fig. 2A). After ~2 months of treatment, this trend slows, as labile compartments (the source of more recently produced virus) decay. If a time-structured tree is constructed only from sequences collected during treatment (Fig. 2B), then a pattern of forward evolution appears. Maximum-likelihood phylogenetic analysis suggests the same pattern, and there are even three internal branches (three separate “novel mutations”) leading exclusively to sequences sampled at the later two timepoints (Fig. 2C). Including sequences sampled before the start of treatment and rooting at the actual infection origin reveals the truth (Fig. 2D): sequences diverge for four months and then the pattern of divergence reverses (i.e., sequences sampled at treatment initiation tend towards rightmost leaves of the tree).

**Figure 2.**
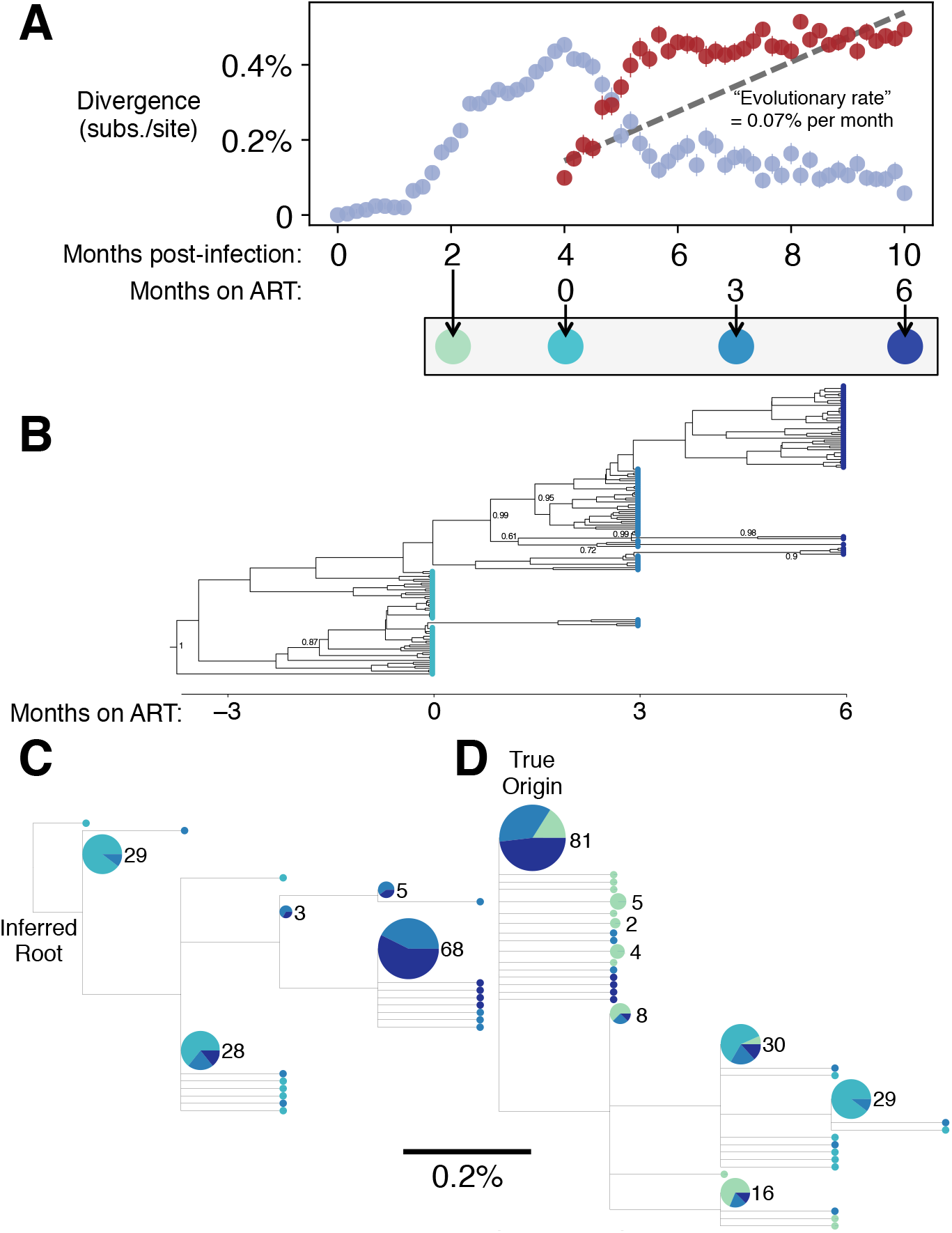
Simulated decay of labile infected cell population creates misleading appearance of viral replication and evolution in first 6 months of treatment. A simulated sample of 50 sequences of length 587 bp is shown at each timepoint (see Methods); these values were chosen for consistency with the number and length of haplotypes presented in Lorenzo-Redondo *et al*.’s phylogenetic analysis. (A) Genetic divergence is measured, following Lorenzo-Redondo *et al*., as the average fraction of sites differing from the most common genotype found at treatment initiation (red symbols). Although simulated treatment halts all viral replication, decay of labile infected cells over the first year of treatment causes the sampled viral population to diverge genetically from this common genotype. If divergence is instead measured from the infection origin (light blue symbols), the true pattern is unmasked: evolution proceeds prior to treatment, but then reverses during early treatment as an increasing number of ancestral reservoir sequences are sampled. (B) A time-structured tree, constructed as in Fig. 1 of Lorenzo-Redondo *et al*. from sequences sampled at and after initiation of ART, creates the misleading appearance of clocklike evolution. (C) A maximum-likelihood tree, constructed and rooted as in Extended Data Fig. 2 of Lorenzo-Redondo *et al*., recapitulates this pattern. A clock-like evolutionary signal is detected (0.07% substitutions per site per month, R^2^=0.54, p<10^−25^). (D) When sequences sampled before initiation of ART are also included in the maximum-likelihood tree and the root is placed at the true origin of infection, no clock-like signal is detected. Legend (A): Bars show standard errors. (B) Posterior clade probabilities > 60% shown. (C – D): Leaf sizes and labels indicate multiplicity of each genotype in the sample; leaves without numbers occur only once; segments show proportion sampled at each time, using the color scheme below panel (A).

We have not aimed to show that 100% of viral replication ceases during suppressive ART; this “absolute negative claim” is neither believable nor strictly necessary. The question relevant to HIV cure research is not whether *any* replication occurs, but whether sufficient replication occurs to fuel viral persistence during ART. Insufficient, or “subcritical” replication may contribute to residual viremia, but does not cause long-term persistence and re-seeding of the latent reservoir. Most importantly, subcritical replication is not a barrier to HIV cure^15^.

What we do claim is that, even in the complete absence of viral replication, misleading evolutionary signatures of high-level replication are expected to appear in the first six months of ART. Adding multiple anatomical compartments or other elaborations to our model could increase realism but would not disturb this basic conclusion. Decay of labile populations confounds evolutionary analysis, and so the observations of Lorenzo-Redondo *et al*. are insufficient evidence for ongoing replication. We thus remain unconvinced that ongoing replication contributes to reservoir stability during ART, and we encourage repetition of these studies using replication-competent viruses from HIV-1-infected individuals after >1 year of suppressive ART.

## Author Contributions

Wrote the manuscript: DISR, ALH, SBL, RFS. Simulations and phylogenetic analyses: DISR, ALH, RFS. Estimation of labile compartments: SBL, RFS.

## Acknowledgements

DISR acknowledges support from amfAR Fellowship # 109511-61-RKRL and National Institutes of Health grant R01GM117591. ALH acknowledges support from National Institutes of Health grant DP50D019851 and Bill & Melinda Gates Foundation award OPP1148627.

## Methods

Simulations used a stochastic model of birth, death, and mutation of infected cells. Upon birth, a productively infected cell gives rise to another infected cell, which may experience mutation. A productively infected cell may transition to any state identified in Fig. 1. Maximum-likelihood trees were constructed using PhyML (HKY85 model; single rate category; estimation of base frequencies, transition/transversion ratio, and proportion invariant sites; best of NNI/SPR). The inferred root was chosen from the earliest timepoint to maximize R^2^ of the root-to-tip regression. The time-structured tree was constructed using BEAST v2.4.4. Details are provided in Supplementary Methods.

**Extended Data Table 1:** Analysis of post-treatment simulated samples based on sequences obtained before treatment by Novitsky *et al*.^14^.

| Number of replicates out of 100<br>that pass test(s)<br>($p < 0.05$) | | Participant Code<sup>14</sup> | | |
| --- | --- | --- | --- | --- |
|  |  | B | H | OU |
| Test 1: Divergence | Forward evolution | 10 | 0 | 36 |
|  | Reverse evolution | 4 | 30 | 0 |
| Test 2: Evolutionary rate | Forward evolution | 24 | 0 | 41 |
|  | Reverse evolution | 2 | 29 | 0 |
| Test 3: Root-to-tip regression | Forward evolution | 20 | 9 | 92 |
|  | Reverse evolution | 56 | 45 | 1 |
| Passes at least one test; not contradicted by other tests | Forward evolution | 22 | 9 | 92 |
|  | Reverse evolution | 42 | 46 | 1 |
| Passes all three tests | Forward evolution | 4 | 0 | 27 |
|  | Reverse evolution | 1 | 27 | 0 |

## Supplementary Methods

### Simulation of intrahost evolution

Intrahost HIV evolution was simulated using a stochastic model of birth, death, and mutation of infected cells. Each simulation starts with a single cell infected with the origin genotype. At a birth event, an actively replicating cell gives rise to another such cell, which has a chance of experiencing mutation at a random site (mutation rate *u* per site, sequenced DNA genome length *L*, HKY model-like nucleotide transition matrix with transition/transversion ratio *κ* and equilibrium frequencies *p_A_, p_C_, P_G_, P_T_*, origin genotype shares these frequencies). An actively replicating cell is “copied” without mutation to one of the three states identified in **Fig. 1** (fast-decay, medium-decay, or stable), at rate *m* for each state. At a death event, an actively replicating cell (per-cell rate *d*), fast-decay cell (rate *d*_1_), or medium-decay cell (rate *d*_2_) is removed from the population. Since the timescales considered are relatively short, cells in the stable reservoir persist throughout the duration of the simulation. When drawing sequence samples, we accounted for the fact that the number of cells in each state vary by several orders of magnitude (while avoiding simulation of a great many cells) by weighting each compartment appropriately.

Birth rate depends on the basic reproductive ratio (*R*_0_), carrying capacity (*C*), current number of actively replicating cells (*x*), and genetic fitness for sequence *i*(*f_i_*). Prior to treatment, the birth rate for sequence *i* is 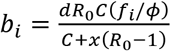, where *ϕ* is the average fitness among actively replicating cells. Note that at carrying capacity (*x* = *C*), for a virus of average fitness (*f_i_* = *ϕ*), the birth rate times the burst size equals the death rate, producing equilibrium. After treatment starts (four months after the start of infection), birth rate is set to zero. If sequence *i* carries no lethal mutations, then its fitness is *f_i_* = exp(*n_p_ s*), where *n_p_* is the number of positively-selected sites mutated in sequence *i* (relative to the origin) and s is the selective advantage; if the sequence carries any lethal mutations, then its fitness is zero. To account for the effect of selection outside the sequenced region (e.g., selection in *env* when *gag* is sequenced), both beneficial and lethal sites may be included in the genome outside the region of length *L*.

#### Parameter values used in all simulations

Basic reproductive ratio *R*_0_ = 6 (peak period, first 20 days of infection), 1.4 (after peak). Death rates *d* = 1/day, *d*_1_ = 0.165/day (half-life of 4.2 days), *d*_2_ = 0.023/day (half-life of 30 days). Mutation rate *u* = 3 × 10^‒5^ per site per generation. Sequenced genome length *L* = 587 bp. Equilibrium nucleotide frequencies *p_A_* = 0.4, *p_C_* = 0.15, *P_G_* = 0.2, *p_T_* = 0.25. Transition/transversion ratio *κ* = 3. Per-cell state transition rate *m* = 0.1/day, with sampling weights 1.82 (fast-decay cells), 4.62 × 10^‒3^ (medium-decay cells), and 5.13 × 10^‒6^ (stable cells) relative to the active cells; these values result in ratios consistent with those determined in **Fig. 1**. During peak period, carrying capacity *C* was 50-fold higher than post-peak. Probability that a site within the sequenced region is lethal when mutated: 70%. Simulations started with a single infected cell and were implemented using discrete steps of 0.25 days; multiple events could occur in a single step.

#### Other parameters

took 50 different sets of values. At least 197 replicates of each parameter set were run. Out of all ∼12,000 runs, 77% were deemed to have a level of viral genetic diversity and divergence at the start of treatment consistent with that reported in studies of untreated patients: We considered average pairwise diversity between 0.092% and 0.285% per site (using the range of four patients with single-founder virus reported in Fig. 6 of Kearney *et al*.^1^ interpolated to 120 days postinfection) and average divergence of 0.009% to 0.578% per site (using the range of estimated evolutionary rates of six patients reported in Fig. 5 of Alizon & Frazer^2^) to be realistic in *gag* and *pol* at four months post-infection.

Results summarized for each of the 50 parameter sets are shown in **Supplementary Table 1**, and detailed results for all runs are shown in **Supplementary Table 2. Fig. 2** of the main text depicts Replicate 91 of Parameter Set 48. A run is judged to have a misleading appearance of forward evolution if it returns a significantly increasing result (p < 0.05) on these three tests:

1. Comparison of average genetic divergence, measured from the most common genotype observed at the start of treatment, between the start of treatment and six months of treatment. (Mann-Whitney U test)
2. Estimate of evolutionary rate computed from linear regression of these divergences over time, using sequences sampled at the start of treatment and at three and six months of treatment. (F-test)
3. Estimate of evolutionary rate, or clock-like phylogenetic signal, computed from root-to-tip regression in the maximum likelihood phylogeny on sequences sampled at the start of treatment and at three and six months of treatment. (F-test)

### Time-structured phylogenies

Time-resolved phylogenies were estimated using BEAST v2.4.4 with a single-compartment version of the model used by Lorenzo-Redondo et al. We used an HKY substitution model with no rate variation, strict molecular clock, and constant size population. More complex models were run but did not substantially improve fit nor change the results. Markov chain Monte Carlo sampling was run for 10^8^ time-steps and convergence was verified using Tracer v1.6.0. The maximum clade credibility tree was chosen from sampled trees for visualization, and posterior clade probabilities >0.5 were annotated. The estimated evolutionary rate was 2.6 ×10^−4^ substitutions per site per month (95% highest posterior density interval 1.0 ×10^−4^ to 4.3 ×10^−4^).

### Simulation of post-treatment decay based on sequences observed during primary infection

As proposed in the Main Text, genetic patterns observed *after* ART has started depend on evolutionary dynamics *before* ART, since pre-ART dynamics determine the relative seeding of cells of varying longevity. The results documented in **Fig. 2** and **Supplementary Tables 1 and 2** rely on the above model of primary infection, which makes assumptions regarding population growth, mutation, and selection. Yet the dynamics of HIV are complex, and a model is always an approximation. Here we describe our approach to sidestep this limitation by using pre-treatment sequence data.

Novitsky *et al*.^3^ longitudinally sampled individuals in Botswana infected with HIV-1 subtype C. Of the cohort of 32, three study participants had samples collected at least three times within the first four months of infection (IDs B, H, and OU). On average there were 13 sequences per time point, available on Genbank as described in the original study. We used only sequences of the *gag* gene (positions 841 to 2,217 of HXB2; *pol* was not sequenced). Sites containing gaps or ambiguous nucleotide calls were ignored. Participant B had 7 sites at which at least two sequences contained nucleotides not present at the first time-point, potential sites under selection. Participants H and OU had 2 and 12 such sites, respectively.

We assumed that the population at each timepoint was reported accurately by the sample. If a sample ***S**_k_* of sequences (some possibly redundant) were observed at time *t_k_*, we assumed that the true population at *t_k_* was composed of these sequences at equal prevalence and no others. We further assumed that the population remained unchanged until *t_k+1_*, at which time it instantaneously changed to match sample ***S***_*k*+1_, based on the sequences observed at that time. We assumed that the first sequence, sampled near the time of seroconversion in all patients, represented the infection origin.

Post-ART population dynamics and simulated sampling followed the model described previously. We assumed that before ART, all viral strains contribute cells to a longer-lived compartment *j* at rate *m_j_*, with values given above. When ART starts, we assume all viral replication is halted, so that seeding stops. All compartments decay at a rate *d_j_* both before and after ART. The expected proportion of the simulated sample observed at time *τ* post-ART that matches a particular sequence present between times *t_k_* and *t_k+1_* is then

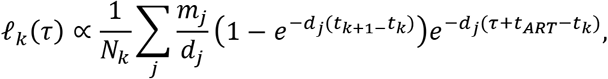

where *N_k_* is the number of sequences sampled at *t_k_* and *t_ART_* is the time at which ART is started (assumed to be 4 months). Note that this formula implicitly assumes that viral load is constant throughout primary infection. Early in infection, viral load starts low, but then peaks briefly above the eventual set-point. As the early timepoints sampled likely included periods both above and below set-point viral load, we chose not to include an explicit viral peak dynamic.

Post-ART samples for each time point (0, 3, and 6 months post-ART) were constructed by random sampling of 50 pre-ART sequences with replacement, weighted by *ℓ_k_*(*τ*). 100 replicates were analyzed by the same tests of forward evolution previously described for the fully simulated data. Results of each replicate are given in Supplementary Table 3; a summary is provided in Extended Data Table 1.

